# Syntrophy between fermentative and purple phototrophic bacteria for carbohydrate-based wastewater treatment

**DOI:** 10.1101/2021.05.13.444055

**Authors:** Marta Cerruti, Guillaume Crosset-Perrotin, Mythili Ananth, Jules L. Rombouts, David G. Weissbrodt

**Affiliations:** Department of Biotechnology, Delft University of Technology, Delft, The Netherlands; Nature’s Principles B.V., Den Haag, The Netherlands

**Keywords:** photoorganoheterotrophs, chemoorganoheterotrophs, syntrophy, mixed-culture fermentation, resource recovery

## Abstract

Fermentative chemoorganoheterotrophic bacteria (FCB) and purple photoorganoheterotrophic bacteria (PPB) are two interesting microbial guilds to process carbohydrate-rich wastewaters. Their interaction has been studied in axenic pure cultures or co-cultures. Little is known about their metabolic interactions in open cultures. We aimed to harness the competitive and syntrophic interactions between PPB and FCB in mixed cultures. We studied the effect of reactor regimes (batch or continuous, CSTR) and illumination modes (continuous irradiation with infrared light, dark, or light/dark diel cycles) on glucose conversions and the ecology of the process. In batch, FCB outcompeted (>80%) PPB, under both dark and infrared light conditions. In CSTR, three FCB populations of *Enterobacteriaceae, Lachnospiraceae* and *Clostridiaceae* were enriched (>70%), while *Rhodobacteraceae* relatives of PPB made 30% of the community. Fermentation products generated from glucose were linked to the dominant FCB. Continuous culturing at a dilution rate of 0.04 h^-1^ helped maintain FCB and PPB in syntrophy: FCB first fermented glucose into volatile fatty acids and alcohols, and PPB grew on fermentation products. Direct supply of carboxylates like acetate under infrared light enriched for PPB (60%) independent of reactor regimes. Ecological engineering of FCB- and PPB-based biorefineries can help treat and valorize carbohydrate-based waste feedstocks.

**GRAPHICAL ABSTRACT:** 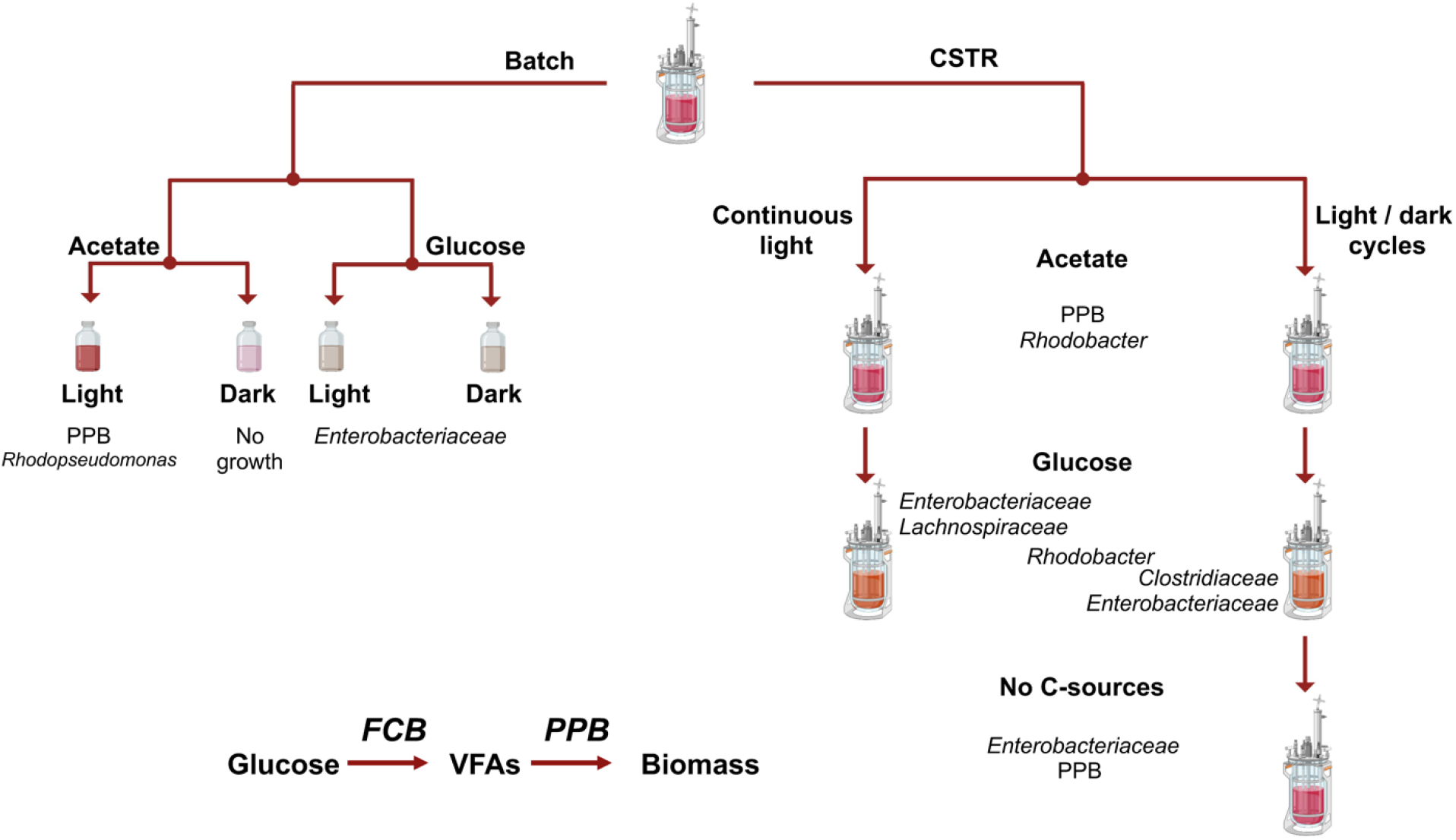

## INTRODUCTION

In the natural environment, bacteria evolved to occupy diverse niches (Steindler et al., 2020). Microorganisms form metabolic associations to grow under nutrient-limiting conditions (Pearman et al., 2008). Anthropogenic environments like biological wastewater treatment plants (WWTPs) involve relatively high concentrations of organics (up to 200 g COD L^-1^) and other nitrogenous and phosphorous nutrients, and complex compositions of carbon sources like cellulose and undigested fibers from food. In agri-food industrial wastewaters, most of organics are derived from simple carbohydrates (*e*.*g*., glucose, fructose or xylose) and polymeric sugars such as starch, cellulose and lignocellulosic derivates (Ghosh et al., 2017).

The wastewater treatment sector transitions to develop bioprocesses that combine nutrient removal with bioproduct valorization. Environmental biotechnologies are studied to treat and valorize nutrient-rich wastewaters by unravelling the biochemical mechanisms that can drive product formation (Kleerebezem & van Loosdrecht, 2007). The efficiency of mixed-culture processes is based on proper selection of microbial guilds needed to metabolize the wastes. Two guilds across chemo- and photoorganoheterotrophic groups are particularly attractive to process carbohydrate- and carboxylate-based feedstocks.

Fermentative chemoorganoheterotrophic bacteria (FCB) degrade carbohydrates through fermentation pathways, producing a spectrum of valuable compounds such as carboxylates (*e*.*g*., volatile fatty acids – VFAs, lactic acid), ethanol and H_2_. Their microbial niches are harnessed in non-axenic mixed-culture fermentation processes to valorize, *e*.*g*., lignocellulosic sugars into targeted fermentation products like lactate or ethanol (Rombouts et al., 2020).

Purple phototrophic bacteria (PPB) are versatile microorganisms, involving a diversity of phototrophic and chemotrophic metabolisms depending on environmental and physiological conditions (**Table 1**). They grow preferentially as photoorganoheterotrophs, harvesting their energy from infrared (IR) light and using a variety of carbon sources from carboxylates to carbohydrates to alcohols (Imam et al., 2011; Okubo et al., 2005; D. Puyol et al., 2017). PPB can also grow as chemotrophs using organic or inorganic substrates (Hunter et al., 2009). As photoheterotrophs, they can achieve high biomass yields up to 1 g COD_X_ g^-1^ COD_S_ (Daniel Puyol & Batstone, 2017), that can be exploited to valorize anabolic products like biomass, single-cell proteins used in animal/human feed, or industrially relevant compounds like carotenoids.

**Table 1.**
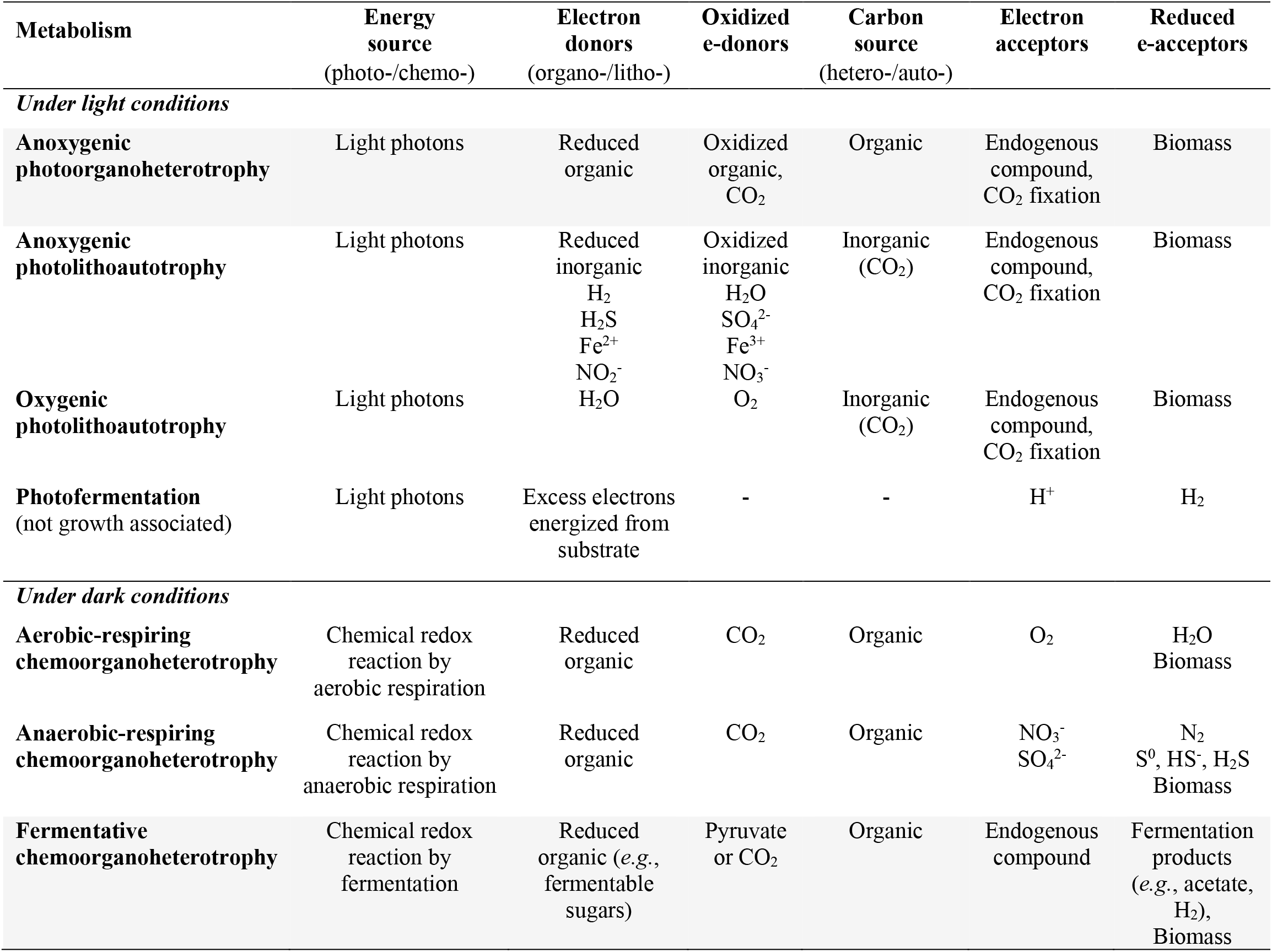
Metabolic versatility of purple phototrophic bacteria (PPB). PPB can grow by combining a diversity of substrates and redox conditions. PPB grow primarily as photoorganoheterotrophs using VFAs as preferential carbon sources. but they are able to grow also on glucose with low growth rates. The metabolisms targeted in this study for the conversion of glucose by combination of fermentation and photoorganoheterotrophy are highlighted in grey.

Further studies evaluated PPB ability to do photofermentation, coupling glucose degradation to H_2_ production. Combinations of fermentative chemoorganoheterotrophy, photoorganoheterotrophy and photofermentation processes have been tested for carbohydrate degradation and conversion. In (*i*) *two-stage dark fermentation and photoorganoheterotrophy*, the process is carried in two separate reactors. In the first unit, pure cultures of FCB degrade carbohydrates into VFAs that are then fed in the second unit to produce PPB biomass in pure cultures (Ghimire et al., 2015). In (*ii*) *single-stage fermentation*, glucose is degraded by pure-cultures of PPB, with longer contact times compared to the two-stages fermentation and the requirement of a pre-sterilized feed stream and axenic environment (Abo-Hashesh & Hallenbeck, 2012). In (*iii*) *single-stage dark and photofermentation*, attempts have been made to co-cultivate defined species of FCB and PPB to simultaneously degrade carbohydrates and convert the produced VFAs to biomass (Rai & Singh, 2016). Despite the advantages of this technique, including lower investment and operation costs compared to the two-staged fermentation, it still requires an axenic environment.

In this context of associating fermentation and photoorganoheterotrophy, one can also wonder whether PPB, due to their metabolic versatility, could be selected in the mixed culture to perform both dark fermentation and photoorganoheterotrophy, or whether FCB would be more efficient on fermentation, prior to supplying fermentation products for the PPB to grow.

Open mixed cultures have the advantage, compared to traditional industrial biotechnology approaches, to not require sterilization of the inflow, therefore substantially reducing the costs (Kleerebezem & van Loosdrecht, 2007). Based on ecological selection principles, microorganisms can be selected to specifically target substrate degradation, bioproduct valorization or both.

Many studies have focused on H_2_ production with FCB and PPB pure cultures or with co-cultures in axenic conditions. Little is known about the community dynamics in open mixed cultures. The interactions between populations and their metabolic interdependency have not been unraveled.

Here, we evaluated the ecological association of PPB and FCB in carbohydrate-fed mixed cultures. We addressed the selection mechanisms, competitive and syntrophic interactions of the two microbial guilds across reactor (batch, chemostat) and IR light irradiance (light, dark, dark/light) regimes using either glucose or acetate as model carbohydrate and carboxylate substrates that harbor the same degree of reduction (4 mol e^-^ C-mol^-1^). With this microbial ecology insights, we tested the possibility to treat carbohydrate-rich wastewaters in a single-stage mixed-culture process assembling FCB and PPB in syntrophy.

## MATERIAL AND METHODS

### Mixed-culture bioreactor systems

To evaluate microbial competition between PPB and fermenters, two mixed-culture bioreactor regimes (batch and continuous culturing), two substrates with the same degree of reduction per carbon mole (acetate and glucose), and three light patterns (continuous illumination, continuous dark, and light / dark switches with a ratio of 12h / 8h) were applied. The reactor performances were measured by quantitative biotechnology endpoints (rates, yields). Microbial selection and community dynamics were tracked by 16S amplicon sequencing. The experimental design is shown in **Table 2**.

**Table 2.**
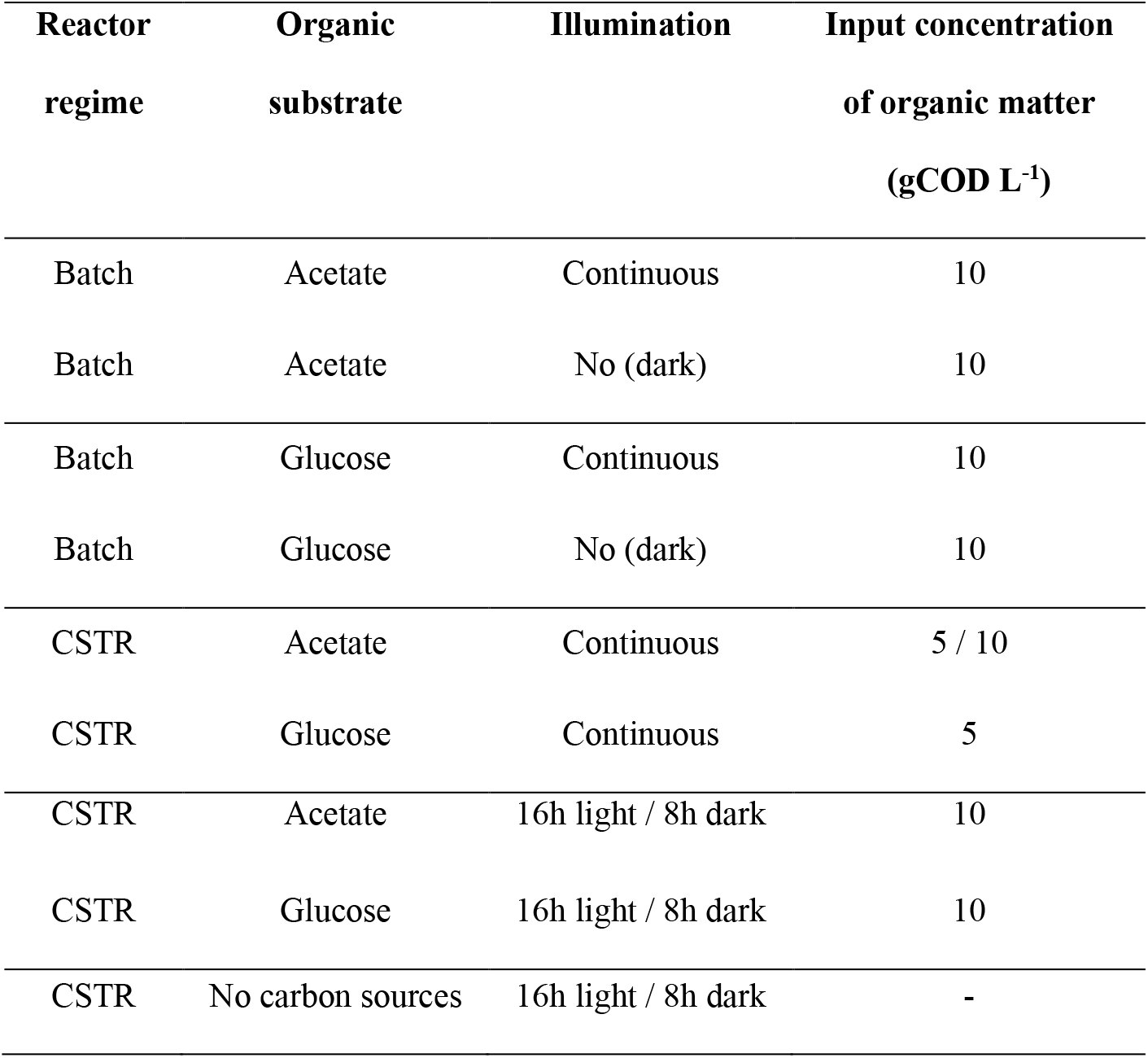
Combinations of operational conditions tested to address selection, competition, and interaction mechanisms between FCB and PPB.

### Compositions of cultivation media

Under all conditions, the cultures were provided with a mineral medium composed of (per L): 0.14 g KH_2_PO_4_, 0.21 g K_2_HPO_4_, 1 g NH_4_Cl, 2 g MgSO_4_·7H_2_O, 1 g CaCl_2_·2H_2_O, 1 g NaCl, and 2 mL of trace elements and 2 mL of vitamins solutions. The stock solution of vitamins was composed of 200 mg thiamine–HCl, 500 mg niacin, 300 mg ρ-amino-benzoic acid, 100 mg pyridoxine–HCl, 50 mg biotin and 50 mg vitamin B12 per liter. The trace elements solution was made of (per L): 1100 mg Na_2_EDTA·2 H_2_O, 2000 mg FeCl_3_·6 H_2_O, 100 mg ZnCl_2_, 64 mg MnSO_4_·H_2_O, 100 mg H_3_BO_3_, 100 mg CoCl_2_·6 H_2_O, 24 mg Na_2_MoO_4_·2 H_2_O, 16 mg CuSO_4_·5 H_2_O, 10 mg NiCl_2_·6 H_2_O and 5 mg NaSeO_3_. The cultivation medium was buffered at pH 7.0 with 4 g L^-1^ 4-(2-hydroxyethyl)-1piperazineethanesulfonic acid (HEPES). Sodium acetate and glucose were used as carbon sources, in a concentration of 5 or 10 g COD L^-1^ depending on experimental conditions (**Table 2**), mimicking concentrations of moderately loaded agri-food wastewaters. Acetate was provided as C_2_H_3_O_2_·3H_2_O (18.28 g L^-1^) and glucose as C_6_H_12_O_6_·H_2_O (9.68 g L^-1^).

### Batches operational conditions

PPB biomass from an in-house enrichment culture grown in a sequencing batch reactor (SBR) (Cerruti et al., 2020) was used to inoculate 100-ml anaerobic bottles. The cultures were flushed with 99% argon gas to maintain anaerobic conditions (Linde, NL, >99% purity). Every batch bottle was duplicated. Half of the batch bottles were exposed to dark. The other half were exposed to IR light (> 700 nm) supplied by a halogen lamp (white light) with a power of 120 W whose wavelengths below 700 nm were filtered out by two filter sheets (Black Perspex 962, Plasticstocktist, UK). Every batch was incubated in a closed shaker (Certomat® BS1, Sartorius Stedim Biotech, Germany) at 30±1°C under dark conditions and at 37±1 °C under light conditions: these differences in temperature were due to an uncontrolled heat derived from the lamps. The flasks were kept under agitation at 170 rpm. The cultures were fed with 10 g COD L^-1^ of either glucose or acetate.

### Continuous culture conditions

A 2.5-L reactor with 2 L of working volume connected to a controller system (In-Control and Power units, Applikon, Netherlands) was used to tune the syntrophy of FCB and PPB under continuous conditions. The pH was set at 7.0 and regulated with NaOH at 0.25 mol L^-1^ and HCl 1 mol L^-1^ and the temperature was maintained at 30°C with a heat exchanger.

The reactor was irradiated from two opposite sides with 2 incandescent lights filtered for wavelengths λ > 700 nm by two sheets (Black Perspex 962, Plasticstocktist, UK) to promote PPB growth. The light intensity on the reactor wall was 270 W m^-2^ and was measured with a pyranometer (CMP3, Kipp & Zonen, The Netherlands).

After start-up under batch mode and PPB enrichment with acetate, the dilution rate was set at 0.04 h^-1^, equal to less than half of the μ_max_ of the PPB representative genus *Rhodopseudomonas* (0.1 h^-1^) (Cerruti et al., 2020).

Four combinations of energy patterns and carbon sources were tested: (*i*) continuous light with acetate, (*ii*) continuous light with glucose, (*iii*) light/ dark (16/8 h) cycles with acetate, and (*iv*) light/ dark (16/8 h) cycles with glucose. Light and dark alternation was controlled with an automatic switch time controlled (GAMMA, The Netherlands).

After 3 months of continuous cultivation, the carbon source feed was stopped, while phosphate and nitrogen sources were maintained for a period of 4 days (ca 4.5 HRTs), in order to evaluate the hypothesis that PPB can grow on fermentation products produced by FCB.

### Mass spectrometry for off-gas analyses

The off-gas of all CSTR was connected to a mass spectrometer (Thermofisher, Prima BT Benchtop MS) which was used to measure the production rates of CO_2_ and H_2_.

### Sampling

All samples were collected in 2 ml Eppendorf tubes, centrifuged at 10000 x *g* for 3 minutes. The biomass pellet was separated from the supernatant and stored at -20°C until further analysis. The supernatant was filtered with 0.2 μm filters (Whatman, US), and stored at -20°C until further analysis. In the batch cultures, the samples were collected every 24 h. During the CSTR, 5 to 9 samples were collected for each condition.

### Biomass determination

Biomass concentrations were measured over the time course of the batch and continuous reactors by spectrophotometry (Biochrom, Libra S11, US) absorbance at 660 nm (A_660_). A calibration curve was established to correlate A_660_ to g volatile suspended solids (gVSS) concentration: c(gVSS L^-1^) = 0.61 A_660_. Biomass gVSS was measured taking samples from the liquid phase, filtering them using 0.2 μm filters (Whatman, 435 USA) and drying the wet filters in a 105 °C stove for 24 h. The filters were then incinerated at 550 °C for 2 h, and weighted.

### Amplicon sequencing

DNA was extracted from 0.5-7 mg biomass using DNeasy Powersoil microbial extraction kit (Qiagen, Hilden, Germany) accordingly to manufacturer’s instructions and stored at -20°C. The compositions of the bacterial communities of the mixed liquors were characterized from the samples collected by V3-V4 16S rRNA gene-based amplicon sequencing as in (Cerruti, Stevens, et al., 2020). The fastq files provided from Novogene were analysed with QIIME2 pipeline (Bolyen et al., 2019).

### Chromatography analysis of substrate conversions and fermentation products

Substrate depletion (acetate and glucose) and fermentation products formation (succinate, propionate, formate, lactate, acetate, butyrate) and were scanned and monitored using high-performance liquid chromatography (HPLC) (Waters, 2707, NL) equipped with an Aminex HPX-87H column (BioRad, USA) column. The eluent used was H_3_PO_4_ (1.5 mmol L^-1^) at a flowrate of 0.6 mL min^-1^ and at a temperature of 60°C. The compounds were detected by refraction index (Waters 2414, US) and UV (210 nm, Waters 2489, US). Ethanol was quantified through gas-chromatography as in (Julius L Rombouts et al., 2019a).

### Wavelength scans of biomass samples to track PPB

Wavelength scan of biomass samples were done using a spectrophotometer (DR3900, Hach, Germany) in order to evaluate the presence of the typical PPB photopigments (carotenoids at 400-500 nm and bacteriochloropyll at ca 850 nm). Absorbance profiles were measured at each wavelength between 320 nm and 999 nm.

### Solver for pathway utilization prediction

To evaluate the fractions of metabolic fermentation and phototrophic pathways, a numerical model was set up which could estimate pathway fractions through minimizing the residual sum squared error (SSQ) using Microsoft Excel 16.48, using the following formula:

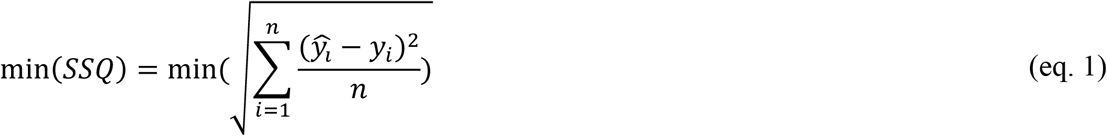

Where *ŷ*_*i*_ is the modelled yield value and y_i_ is the measured yield value, and *n* is the amount of metabolic compounds that are considered. The estimated yields were calculated using different fractions of metabolic pathways, which could be varied by the solver to minimize the SSQ (**Table 3**). This SSQ value was minimized using the Generalized Reduced Gradient (GRG) Nonlinear solver available in Excel. The pathways that were used varied per enrichment, depending on the microbial community structure, assuming only the activity of relevant catabolic and anabolic pathways. To simulate the batch enrichments the equations 1.1-1.8 were used. To simulate the CSTRs, reactions 1.9-.10 were also added (Table 3). The result of the pathway modelling was used to create **Figure 7**, visualizing in a semi-quantitative way the flux distribution of the different pathways.

**Table 3.** Putative reactions included in the numerical model of metabolic reactions along FCB and PPB pathways. The reactions 1.1-1.8 were used to evaluate the contribution of each pathway under batch conditions. The reactions 1.9 and 1.10 were included in the simulation of the CSTRs.

| Microbial guilds | Pathways | Reactions |
| --- | --- | --- |
| **FCB** | Glucose → Biomass + CO<sub>2</sub> | (1.1) |
| | Glucose → Ethanol + CO<sub>2</sub> | (1.2) |
| | Glucose + H<sub>2</sub>O → Acetate + Ethanol + H<sup>+</sup> + H<sub>2</sub> + CO<sub>2</sub> | (1.3) |
| | Glucose → Butyrate + CO<sub>2</sub> + H<sub>2</sub>O | (1.4) |
| | Glucose → Succinate | (1.5) |
| | Glucose → Lactate + H<sup>+</sup> | (1.6) |
| | Lactate → Acetate + Propionate + CO<sub>2</sub> + H<sub>2</sub>O | (1.7) |
| **FCB + PPB** | Formate → CO<sub>2</sub> + H<sub>2</sub> | (1.8) |
| **PPB** | Acetate → Biomass + CO<sub>2</sub> | (1.9) |
| | CO<sub>2</sub> + H<sub>2</sub>O → Biomass + H<sub>2</sub> | (1.10) |

## RESULTS & DISCUSSION

The batch and continuous reactor experiments revealed key insights on the microbial community engineering principles governing the selection, competition and interaction between FCB and PPB in function of substrates, light patterns, and reactor regimes.

### Batch conditions

#### In mixed-culture batch mode, PPB were selected on acetate and IR light

Under batch conditions, the acetate-fed cultures grew only when subjected to light, reaching a biomass concentration of 1 gVSS L^-1^ (µ= 0.0411 ± 0.0007 h^-1^), with a yield of biomass on substrate of 0.60 ± 0.09 C-mmol_X_ C-mmol_S_^-1^. Little growth was reported in the six days of cultivation for the acetate cultures in the dark, with a biomass specific growth rate (µ) of 0.006 h^-1^.

The inocula of the batches originated from a PPB in-house enrichment culture operated as SBR in parallel to the experiments. The inoculum seeded in the acetate-fed batches presented a relative abundance above 60% (vs. total amplicon sequencing read counts) of the purple non-sulfur genus *Rhodopseudomonas*. In the acetate-fed batches, *Rhodopseudomonas* sp. reached as high as 80% of the total community after 6 days of cultivation under IR light. Despite the low growth, chemoorganoheterotrophs of the family of the *Moraxellaceae* showed the highest relative abundance (80%) in the dark (**Figure 1A**).

**Figure 1.**
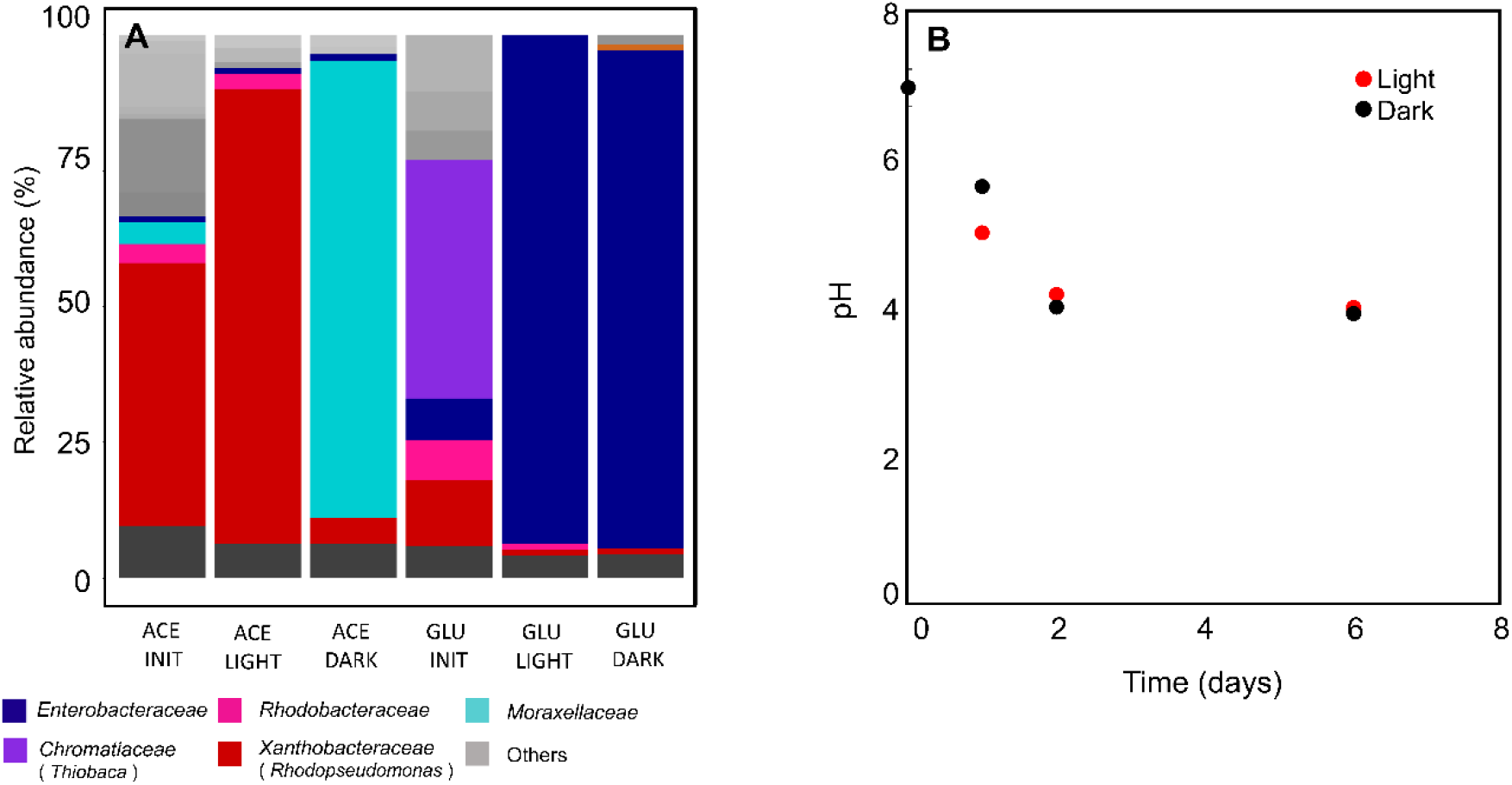
**A:** Microbial community composition in batches. PPB are dominant at the beginning of the cultivation (ACE INIT and GLU INIT). After 6 days of cultivation under acetate feed, PPB were dominant under light conditions (ACE LIGHT). Under acetate-dark conditions there was little grow from the genus *Moraxellaceae*. Under glucose feed, independently of the light conditions, the genus Enterobacteriaceae was dominant. **B:** pH evolution in the glucose-fed batches. Both under light and under dark conditions the pH dropped to 4 in the first 2 days of incubation.

Acetate is one of the preferred substrates for PPB (Blankenship et al., 1995). It is assimilated into biomass by many PPB species, through the tricarboxylic acids cycle (Pechter et al., 2016). Acetate and IR light selected for *Rhodopseudomonas* under acetate/light conditions. Among the PPB, *Rhodopseudomonas* has the highest growth rate on acetate (µ_max_ = 0.15 h^-1^) (Cerruti, Ouboter, et al., 2020) and therefore was predominant at the end of the batch cultivation. The growth rate in the acetate-fed cultures under dark conditions was 7 times lower than under light conditions, and the biomass concentration after 6 days was 270 times lower than under light conditions. Under dark conditions, acetate can only be assimilated in presence of suitable electron acceptors (such as S^0^ or SO_4_^2-^) in order to produce ATP for growth (Madigan et al., 1982). In the batches, little external electron acceptor was present in the medium (3 mmol SO_4_^2-^ L^-1^). This can explain the very low biomass growth under dark conditions compared to the light one.

#### In mixed-culture batch mode, FCB were selected on glucose

In the glucose-fed batches a higher biomass concentration was reached in the dark (1 gVSS L^-1^) than in the light (0.4 gVSS L^-1^), with a specific µ of 0.070 ± 0.001 h^-1^ and 0.020 ± 0.002 h^-1^ respectively. The inoculum of glucose-fed batches was dominated by the purple sulfur genus *Thiobaca*, due to microbial community variations in the parent reactor. In the glucose batch cultivation, the genus of *Enterobacter* was predominant, above 90% of the total sequencing reads (**Figure 1A**), both under light and dark conditions. The pH was not controlled in batch cultivations: in the glucose-fed batches pH dropped to 4 within 48 h, despite the presence of the HEPES buffer (**Figure 1B**).

The *Enterobacteriaceae* family is known to be predominant in anaerobic environment in presence of non-limiting concentration of monomeric carbohydrates, such as glucose (de Vrije & Claassen, 2003) (de Vrije & Claassen, 2003). Rombouts et al., 2019 reported an enrichment of 75% of the genus *Enterobacter* in a glucose-fed SBR, in nutrient, temperature and pH conditions similar to the one here presented.

#### Ethanol was the main fermentation product under glucose-fed batch conditions

Under acetate-fed batch conditions in the light, acetate was fully depleted after 6 days of incubation. Under glucose-fed batch conditions the biomass yield over glucose were 0.052 ± 0.002 C-mmol_X_ C-mmol_S_^-1^ and 0.134 ± 0.011 C-mmol_X_ C-mmol_S_^-1^ under light and under dark conditions respectively. After six days of cultivation, in the glucose-fed batches in the dark, ethanol was the main fermentation product, with a yield of 0.20 ± 0.10 C-mmol_et_ C-mmol_S_^-1^, followed by succinate (0.07 ± 0.03 C-mmol_succ_ C-mmol_S_^-1^) and acetate (0.03 ± 0.01 C-mmol_ace_ C-mmol_S_-1). In the glucose-fed batches under light conditions, after 6 days of cultivation 30% of the initial glucose was still present in the reaction broth, and carbon balances could not be properly closed (>90% recovery). Ethanol was the main fermentation product (0.10 ± 0.07 C-mmol_et_ C-mmol_S_^-1^), followed by lactate (0.05 C-mmol_lac_ C-mmol_S_^-1^) and succinate (0.04 ± 0.02 C-mmol_succ_ C-mmol_S_^-1^).

A solver enabled to evaluate the theoretical contribution of 8 putative fermentation pathways to the overall carbon and electron balances. Under batch conditions the phototrophic pathways were not included in the simulation, as PPB relative abundance was very low as measured by amplicon sequencing (∼1%) (**Figure 1A**). The predicted contribution of the singular pathways to the overall yields of the batches under light and dark conditions were comparable. Under batch conditions, the dominant metabolic pathway was identified as homo-ethanol production form glucose (**Table 3**, Eq. 1.2). This catabolism is associated with the fermentative *Enterobacteriaceae* (Jang et al., 2017).

Wavelength scan is commonly used as an indirect measure for the presence of PPB in the cultures (Fiedor et al., 2016), as photopigments absorb photons with specific wavelengths. Absorbance peaks around 400-500 nm represent the carotenoids series, and the peaks around 850 nm the bacteriochlorophyll a (Stomp et al., 2007). In the acetate-based batches in the light the aforementioned peaks were clearly detected at 450 nm and 800-850 nm. In the acetate-fed batches in the dark, almost no growth was reported in the 6 days incubation and peak for photopigments was detected at 800 and 850 nm. In the glucose-fed batches, both dark and light, no peaks were observed (**Figure 2A/B**). These results suggested the selective growth of PPB in illuminated acetate batches, and of non-phototrophic organisms in the glucose-batches.

**Figure 2.**
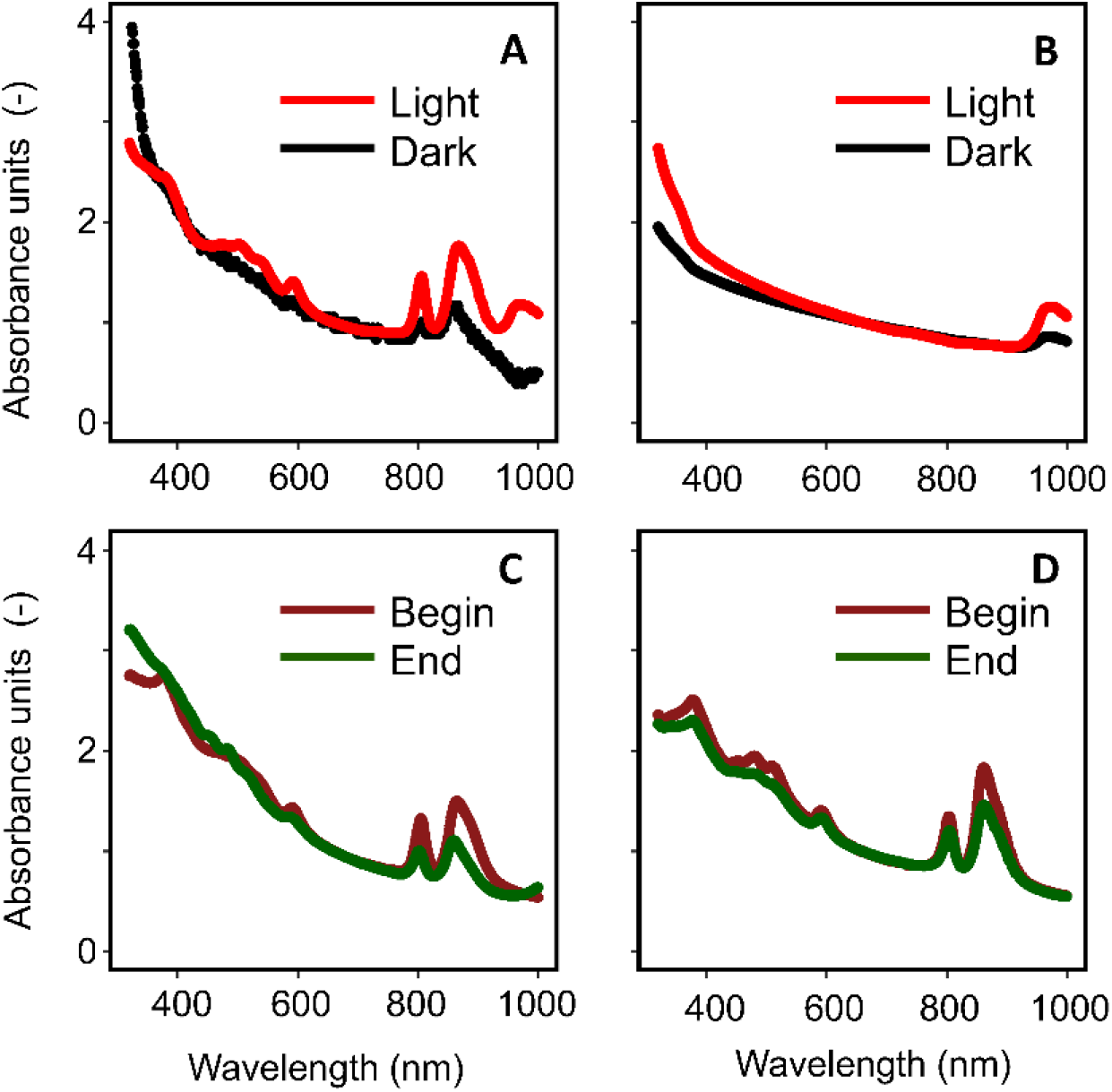
Wavelength scan from 300 to 1000 nm. Absorbance units normalized for the biomass concentration (as A.U. at 660nm). **A:** acetate-fed batches. The typical PPB peaks (at ca 850 nm) were clearly visible under light conditions. **B:** glucose-fed batches. No peak was present. **C:** continuous light glucose-fed CSTR. The presence of PPB was confirmed by the presence of their typical peaks. **D:** dark / light glucose-fed CSTR. The presence of PPB was confirmed by the presence of their typical peaks.

#### Enterobacteriaceae were enriched due to higher growth rates

The maximum biomass specific growth rate (µ_max_) is a key factor driving selection mechanisms in substrate-rich environments (Winkler et al., 2017), if no significant intermediate storage of substrate is displayed. FCB-like species of the genus *Enterobacter* present a µ_max_ of 0.45 – 0.80 h^-1^ in mineral medium conditions with low sugar concentrations (<10 g_COD_ L^-1^) (Füchslin et al., 2012; Rombouts et al., 2019). When fermenting carbohydrates like glucose, *Rhodopseudomonas capsulata* grows at a µ_max_ of 0.014 h ^-1^ (Conrad & Schlegel, 1978). In batches, where the substrate is not limiting, the organisms with higher uptakes rates will be selected above organisms with high biomass yields (Rombouts et al 2019). According to the Herbert-Pirt relation, when the substrate uptake is directly coupled with growth, the organisms with the highest growth rate dominate. Using these studies, the genus *Enterobacter* is estimated to have a µ_max_ about 32 times higher than *Rhodopseudomonas* under glucose feed, making it successful in the competition for glucose in batch.

Under batch conditions, both in the light and in the dark, the family of the *Enterobacteriaceae* was enriched (>90%). The species belonging to the family of the *Enterobacteriaceae* ferment sugars through either the Embden-Meyerhof pathway or the pentose phosphate pathway, with a net production of ethanol (Jang et al., 2017). Ethanol can be used as substrate for growth by PPB, but with a strict pH regulation (pH = 7) (Inui et al., 1995).

Due to lactate production, which is a major byproduct of ethanol fermentation (Converti & Perego, 2002), and insufficient pH buffer, the pH dropped to 4 within the first 2 days of cultivation, as the pK_a_ of lactic acid is 3.8 (**Figure 1B**). Fermentative bacteria can still grow at low pH (pH < 5.5) (Tsuji et al., 1982). PPB grow instead in a pH range between 6.5 and 9.0, with an optimum at 7.0 (Pfennig, 1977). The low pH (pH = 4) resulting from the glucose-fed batch experiments could have prevented the PPB to uptake and grow on the alcohols and VFAs produced by *Enterobacter*, leading to a low relative abundance (∼ 1%) in the glucose-fed batches. We postulate that with a strict pH control to 7, PPB will be able to grow on fermentation products of *Enterobacteriaceae*.

The genus *Enterobacter* can grow in a range of temperature between 20 and 45° C (Gill & Suisted, 1978), with an optimum growth temperature at 40° C (Tanisho, 1998). The temperature differences here reported for glucose-fed batches under dark and light (ca. 7° C difference) should not have affected its specific growth rates. On the other hand, light has been used as a control method for fermentative organism proliferation (D’Ercole et al., 2016; Gwynne & Gallagher, 2018). As IR light was provided to the illuminated cultures, we postulate its inhibitory action on the *Enterobacteriaceae* family, resulting in lower growth rates (3.5 times lower) compared to the dark cultures. How this inhibition by photons actually works on cellular and molecular level remains to be settled by future work.

### Continuous culture

Under continuous-flow regime, four different light and substrate conditions were imposed (1-acetate under light, 2-acetate under light / dark cycles, 3-glucose under light, and 4-glucose under light and dark cycles). The pH was controlled at 7.0 to favor the enrichment of PPB.

#### CO_2_ production rate increased with an increased biomass growth

The CO_2_ partial pressure in the off-gas of the CSTR was monitored to evaluate the growth state of the biomass. For PPB, CO_2_ production is a side product of acetate assimilation (Sadaie et al., 1997). In acetate-fed CSTRs, CO_2_ production was constant under continuous illumination (0.0111 ± 0.0004 C-mmol_CO2_ C-mmol_S_^-1^) (**Figure 3A**). When dark / light cycles were applied, the CO_2_ production decreased to 0 C-mmol h^-1^ during the 8 h dark periods and increased again to 0.3 C-mmol h^-1^ during the 16 h light phases (**Figure 3B**). The CO_2_ production during the light phases indicated that acetate was continuously assimilated, whereas the sudden decrease during the dark periods (1/3 less within the first hour of dark) indicated that the metabolic activity was switched off. An oscillation in the CO_2_ production rate was already observed in *Rhodopseudomonas* pure cultures under CSTR (Cerruti et al., 2020). In both cases, when the substrate was provided in excess, PPB grew with a growth rate close to μ_max_ (0.1 h^-1^) during the light periods. In the dark, PPB halted their metabolic activity.

**Figure 3.**
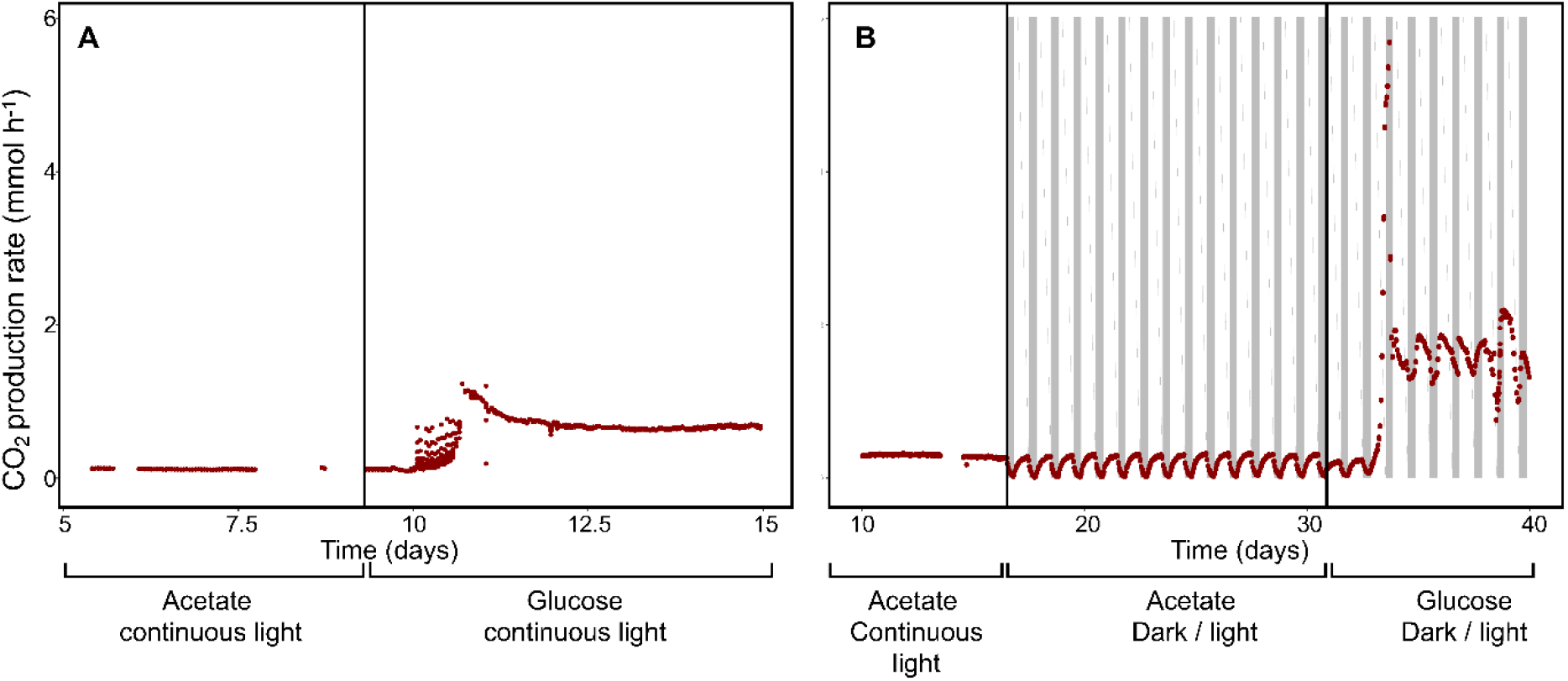
**A:** CO_2_ production rates under acetate-fed and glucose-fed CSTR continuous light conditions. An increase of ca 1 mmol h-1 was observed 3 days after the switch to glucose feed, and then stabilized. **B:** CO_2_ production rates under acetate continuous light, acetate dark / light and glucose dark / light conditions. Under acetate continuous illumination CO_2_ was produced at constant rate. Under acetate dark / light CO_2_ was produced only during the light periods. Under glucose dark / light conditions, a peak in production rate was observed after 3 days from the switch to the glucose-feed. The production rates increased during dark periods and decreased during light periods.

Both under continuous illumination and dark/light cycles, a change in CO_2_ production was observed 24-48 h after the switch from acetate feed to glucose feed. In the CSTR under continuous illumination, a peak of CO_2_ production reached 1.2 C-mmol h^-1^ 24 h (ca 1 HRT) after the switch to the glucose feed and stabilized at 0.6 C-mmol h^-1^ 2 days later (**Figure 3A**). Under light / dark cycles a peak of CO_2_ production (up to 5.6 C-mmol h^-1^) was observed after two full cycles (3.8 HRTs from the glucose switch). After 8 HRTs from the switch to the glucose feed, the CO_2_ production rate presented oscillations between light and dark periods. The CO_2_ production rate increased of 0.5 C-mmol h^-1^ at the beginning of the light phase and it decreased of 0.4 C-mmol h^- 1^ in every dark period (**Figure 3B**). This behavior was observed for 8 HRTs, until the pH dropped to 2 due to a technical failure.

#### Ethanol was the main fermentation product also in glucose-fed CSTR

Under glucose-fed CSTR conditions, the biomass yield over glucose was 0.22 ± 0.02 C-mmol_X_ C-mmol_S_^-1^ and 0.45 ± 0.12 C-mmolX C-mmol_S_^-1^ under continuous illumination and light / dark cycles respectively. Ethanol was the main measured fermentation product (0.18 ± 0.06 and 0.11 ± 0.10 C-mmol_et_ C-mmol_S_^-1^ under continuous light and dark / light cycles respectively). In the CSTR under continuous illumination also acetate was measured (0.21 ± 0.15 C-mmol_ace_ C-mmol_S_^-1^) and butyrate (0.06 ± 0.05 C-mmol_but_ C-mmol_S_^-1^). In the CSTR under dark / light cycles, formate (0.11 ± 0.01 C-mmol_form_ C-mmol_S_^-1^) and butyrate (0.10 ± 0.01 C-mmol_but_ C-mmol_S_^-1^) were measured (**Figure 4**).

**Figure 4.**
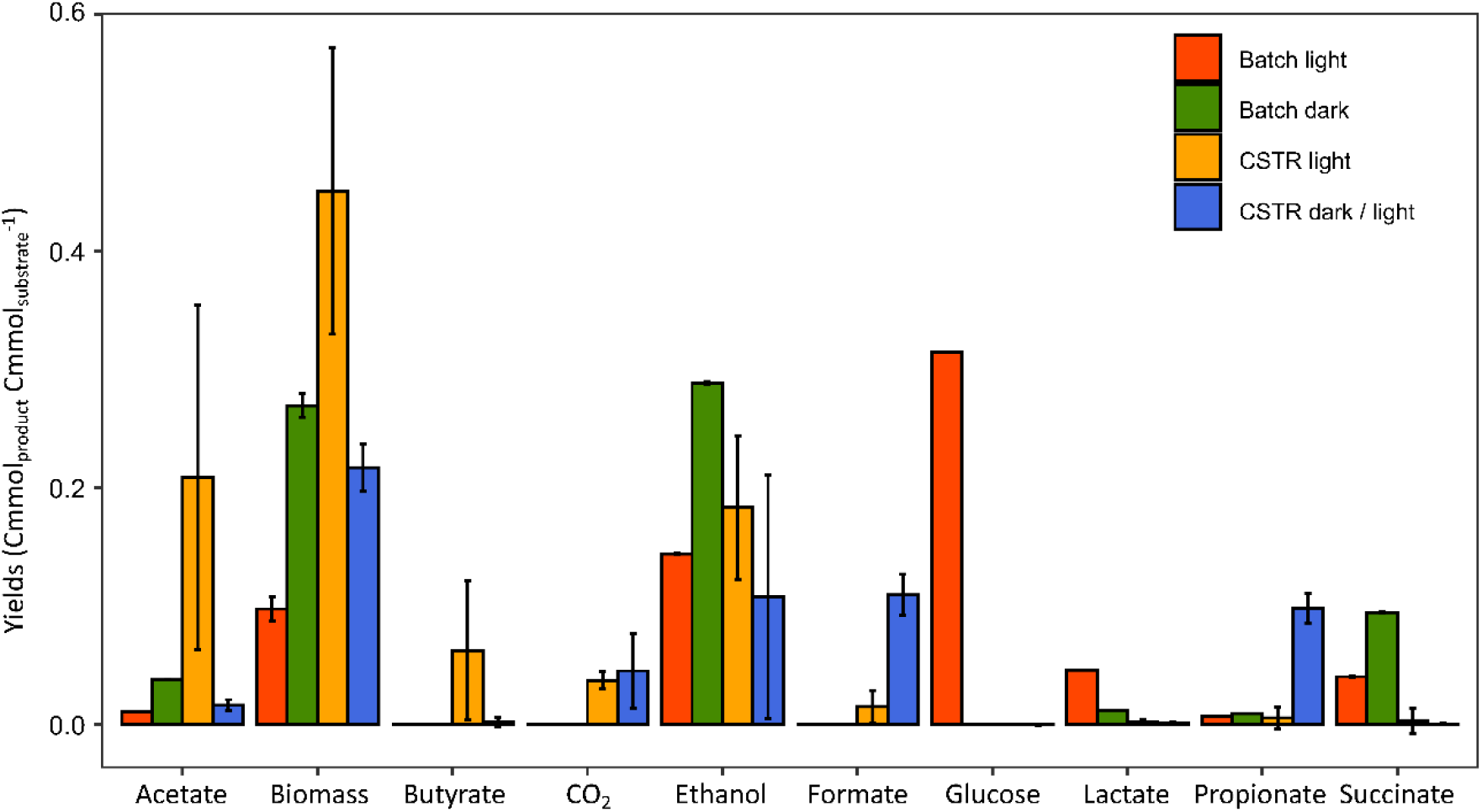
Yields of the glucose-fed cultures. Under batch conditions, the main fermentation products were ethanol and succinate. Under light conditions, glucose was not fully depleted. Under CSTR conditions, the main fermentation products were ethanol, formate and acetate.

Under CSTR conditions, photoautotrophic CO_2_ fixation (photosynthesis) and photoheterotrophic acetate assimilation were added to the solver matrix, to evaluate the contribution of PPB to the total carbon and energy balances. Under all CSTR conditions the major fermentation pathway involved the glucose fermentation through the acetyl-CoA pathway (**Figure 5**) (83% and 55% for continuous illumination and dark / light cycles respectively). The solver estimated a high contribution of the photoautotrophic pathway in the CSTR under continuous illumination, and of the photoheterotrophic pathway under dark / light cycles.

**Figure 5.**
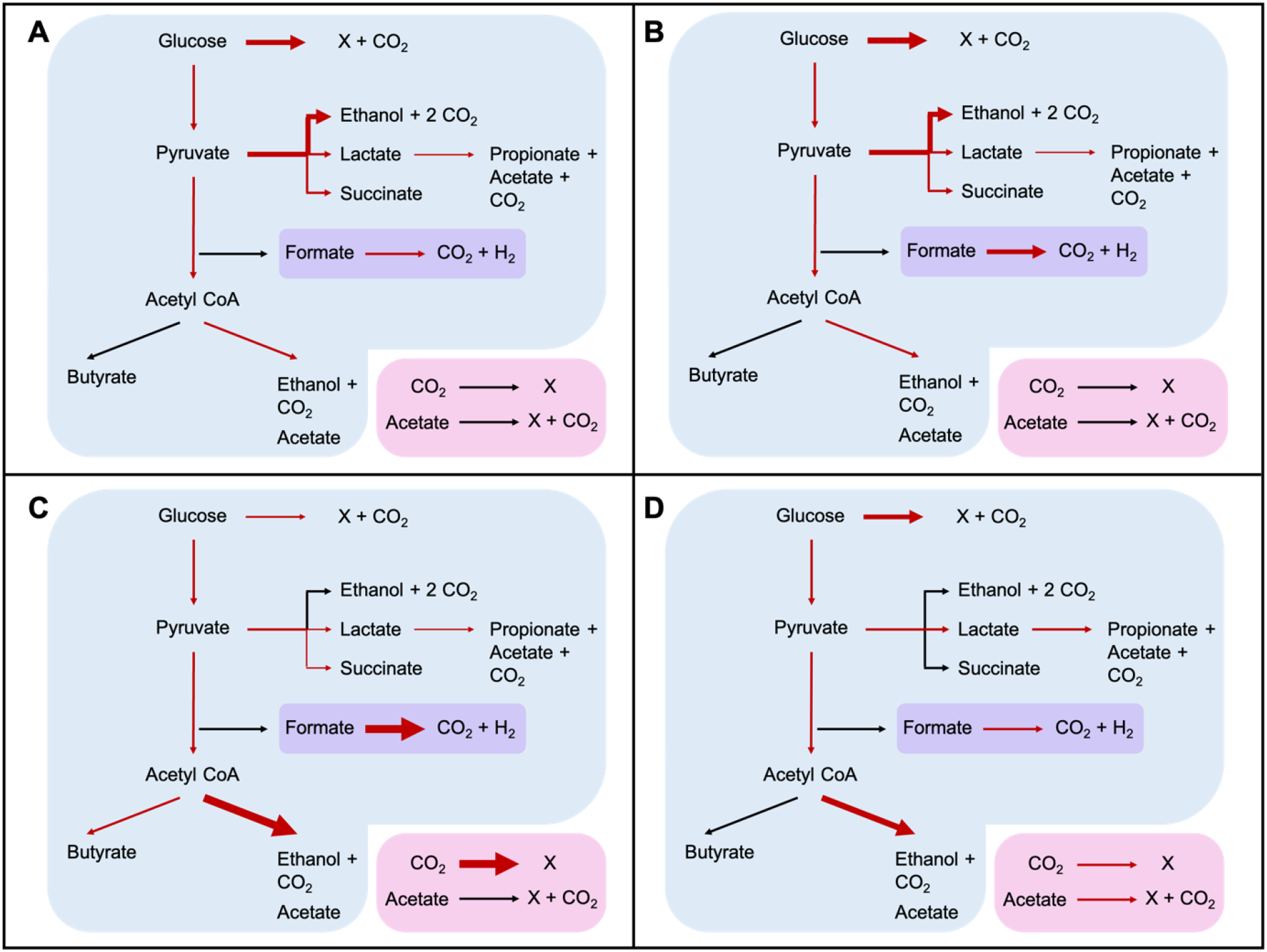
Pathways involved under the different reactor regimes. The thickness of the arrows represents the predicted yield toward a specific metabolic route, based on the calculations of the solver. The reactions with the blue background are performed by FCB. The reaction with the purple background is performed by both FCB and PPB. The reactions with the pink background are performed by PPB. **A:** batch conditions with continuous illumination. **B:** batch conditions in the dark. **C:** CSTR under continuous illumination. **D:** CSTR conditions under light / dark cycles. Under batch conditions the pathways used were mainly directed toward biomass and ethanol production. Under CSTR ethanol was the main fermentation product, but the production of propionate, succinate and lactate was predicted. The phototrophic pathways were not included for the batch conditions but were included in the carbon and electron balance of the CSTRs.

#### PPB were enriched in the acetate-fed CSTR

Under continuous illumination and with acetate as carbon and electron source PPB formed the dominant guild (on average 68 ± 21 %) (**Figure 6**). The genus *Rhodopseudomonas* was predominant at the beginning of the CSTR (65%), while its relative abundance decreased to 40% after one week of cultivation (8.5 HRT). Simultaneously, the genus *Rhodobacter* got enriched (from 13 to 23%). A further enrichment of *Rhodobacter* was observed once the light/dark cycles were applied, reaching around 65% of the total population after 15.6 HRTs, while *Rhodopseudomonas* abundance was stable below 10%.

**Figure 6.**
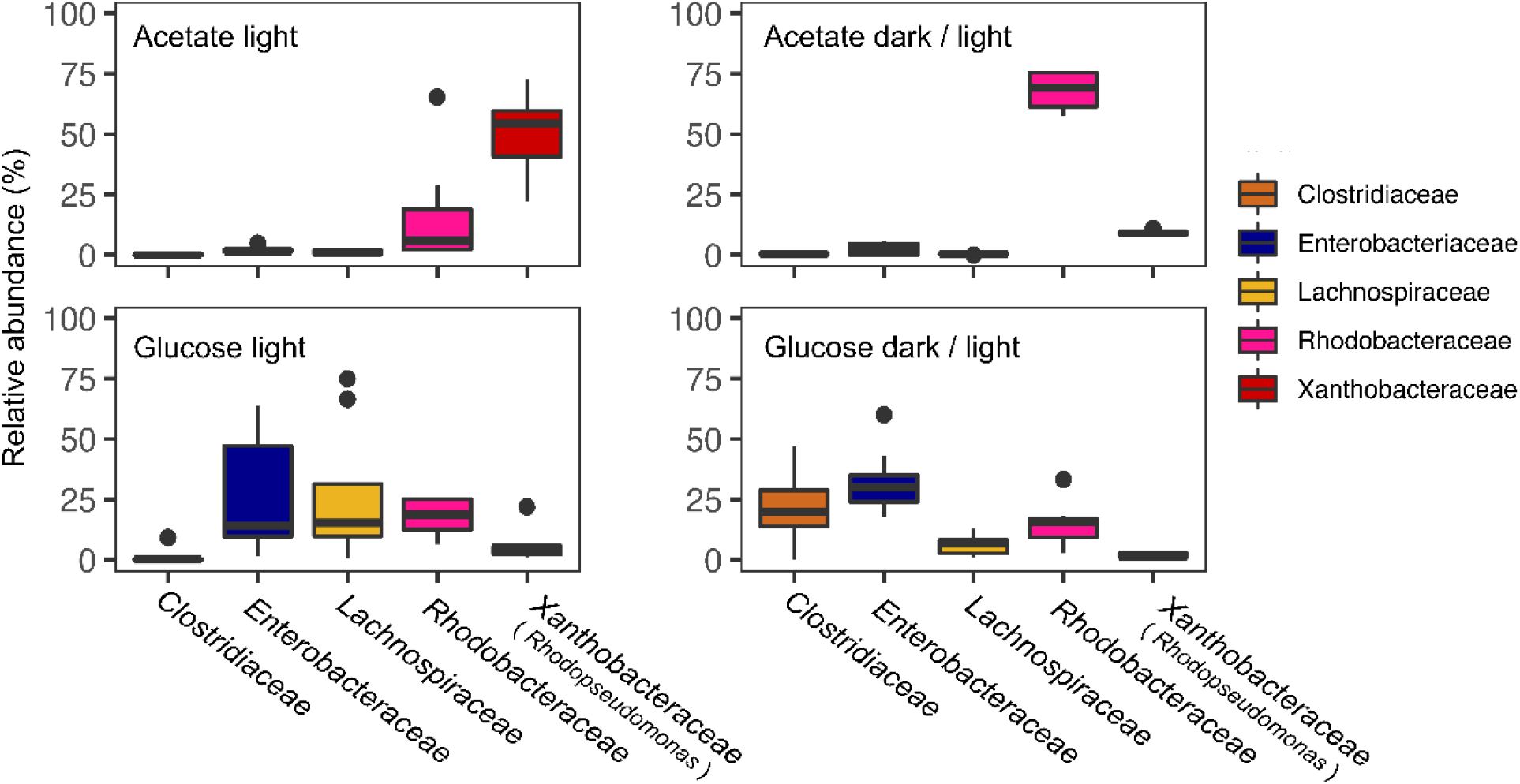
Relative abundance under CSTR conditions. Under acetate conditions, PPB were dominant community, with a shift from *Rhodopseudomonas* under continuous illumination to *Rhodobacter* under light / dark conditions. Under glucose feed, the FCB were dominant, especially the genera *Enterobacteriaceae* and *Lachnospiraceae*.

In the acetate-fed CSTR under continuous light the biomass concentration was not constant. Once the dark / light cycles were applied, the biomass production followed the irradiation pattern. The biomass reached a concentration of 44.75 ± 1.24 C-mmol L^-1^ after the light periods, and it decreased to 37.58 ± 0.09 C-mmol L^-1^ after the dark periods. Under anaerobic conditions, PPB do not grow in the dark in absence of an external electron donor (Madigan et al., 1982; Yen & Marrs, 1977), but growth is restored once the light is present (Cerruti et al., 2020; Zhi et al., 2019). The biomass concentration increased without stabilizing in the light periods, as in (Cerruti et al., 2020). Acetate was not consumed during the dark phases, and its concentration increased due to continuous supply in the CSTR. In a chemostat, the growth rate of the microbial community is defined by the dilution rate and the concentration of the limiting compound, if no biofilm is formed (Kuenen, 2019). An increase in biomass concentration during the illumination times implied that the growth rate of the microorganisms (0.1 h^-1^) was higher than the dilution rate (0.04 h^-1^), as a consequence of high residual acetate concentration. During the dark cycles, acetate consumption stopped, acetate accumulated, to be used in light cycles by the PPB organisms. This a higher growth rate than the applied dilution rate, which fits with standard Monod kinetics for growth.

#### FCB were enriched in the glucose-fed CSTR

In the CSTR, FCB accounted for more of the 50% of the sequences under glucose feed, regardless the light condition. In CSTR, light and dark cycles had an impact on the selection process of the FCB. Under continuous illumination, the family of *Lachnospiraceae* was predominant (around 30 % of the total community, with a peak at 75%), followed by the family of *Enterobacteriaceae* (15%) (**Figure 6**). Under light/dark cycles *Enterobacteriaceae* was the most predominant (32 ± 17 %), along with *Clostridiaceae* (18 ± 15 %).

Among the PPB guild, *Rhodobacter* was the only PPB to significantly persist in the glucose-fed CSTR, with an average relative abundance of 19.5 ± 14 % (**Figure 6**) independently of the light conditions. The persistence of PPB in the glucose-fed CSTRs was further proven by the wavelength scan. Peaks for the bacteriochlorphyll and the carotenoids were identified at 400-450, 800 and 850 nm for both the continuous illumination and the light / dark cycles (**Figure 2C/D**).

#### The microbial community defined the fermentation products production

The families of *Clostridiaceae, Enterobacteri*aceae and *Lachnospiraceae* are able to ferment glucose. They compete for glucose through different fermentation pathways (Grimmler et al., 2011; Horiuchi et al., 2002). *Lachnospiraceae* and *Clostridiaceae* are phylogenetically and morphologically heterogeneous families belonging to the phylum of *Firmicutes* (De Vos et al., 2005). Glucose fermentation through the acetyl-CoA pathway is typical of *Clostridiaceae* (Aristilde et al., 2015). In human gut microbiota, they contribute to sugar fermentation to lactate and short-chain fatty acids production (Venegas et al., 2019). Butyrate was produced only in the CSTR under continuous illumination (0.06 ± 0.05 C-mmol C-mmol_S_^-1^). The *Lachnospiraceae* and *Clostridiaceae* families ferment glucose into primarily butyrate, acetate and ethanol (Rombouts et al., 2019b; Temudo et al., 2008; Valk et al., 2018). Butyrate is produced through the butyril-CoA dehydrogenase gene, responsible for the conversion of crotonyl-CoA to butyryl-CoA and the butyryl-CoA:acetyl-CoA transferase (Venegas et al., 2019). According to the NCBI database (NCBI, 2021), the gene encoding for this enzyme is present in the phylum of *Firmicutes* that comprises *Clostridiaceae* and *Lachnospiraceae* relatives, but not in the proteobacterial family of the *Enterobacteriaceae*, explaining the absence of butyrate in the glucose-fed batches.

Under dark / light CSTR conditions, the relatively high production of propionate (0.10 ± 0.01 C-mmol C-mmol_S_^-1^, 17 times higher than under continuous illumination), can be linked to the abundance of *Clostridiaceae*, which are propionate producers (Johns, 1952). *Clostridium* species can present a µ_max_ of 0.25 h^-1^ (Gomez-Flores et al., 2015) with primary fermentation products including ethanol, butyrate, acetate and propionate (Huang et al., 1986; Lamed et al., 1988). *Clostridium beijerincki* has been reported to decrease the H_2_ production when exposed to light intensities above 200 W m^-2^, with a shift from a preferential production of butyric acid to acetic acid (Zagrodnik & Laniecki, 2016). The continuous illumination under CSTR conditions might have resulted in an inhibition of *Clostridiaceae*, with an enrichment of other *Firmicutes* like *Lachnospiraceae*, through an unknown mechanism.

Formate and propionate production was 10 times higher under light/ dark conditions compared to continuous illumination. Acetate production was instead 12 times lower under dark / light cycles compared to the continuous illumination (**Figure 4**). This was probably linked to a higher abundance family of the *Enterobacteriaceae* under dark / light cycles compared to the continuous illumination (31% and 18% respectively) (**Figure 6**). *Enterobacteriaceae* present a µ_max_ 0.5-1 h^-1^ (Khanna et al., 2012; Martens et al., 1999) and are known to ferment sugars to primarily ethanol, with lactate and acetate as major byproduct during ethanol fermentation (Converti & Perego, 2002).

#### PPB grew on FCB fermentation products

After 3 months of CSTR operation, to test the hypothesis that PPB grow on the fermentation products of FCB, the glucose feed was stopped while nitrogen and phosphate were still provided. Along the 4 days of operation, the relative abundance of the PPB shifted from ca 10 to 50% (**Figure 7A**). The biomass concentration sharply decreased already 2 h after stopping the glucose feed, stabilizing at a concentration of 15.7 ± 2 C-mmol_X_ L^-1^. Butyrate, ethanol, formate, lactate, propionate and succinate were washed out of the system at the imposed dilution rate (0.04 h^-1^). Acetate was depleted in 50 h, 4 times faster than the theoretical washout rate (**Figure 7B**). In absence of glucose as carbon source, the highly active fermentative organisms were not growing. The increase in the PPB relative abundance proved the interaction with FCB, showing that PPB utilize VFAs as preferred substrate and can selectively grow on FCB fermentation products.

**Figure 7.**
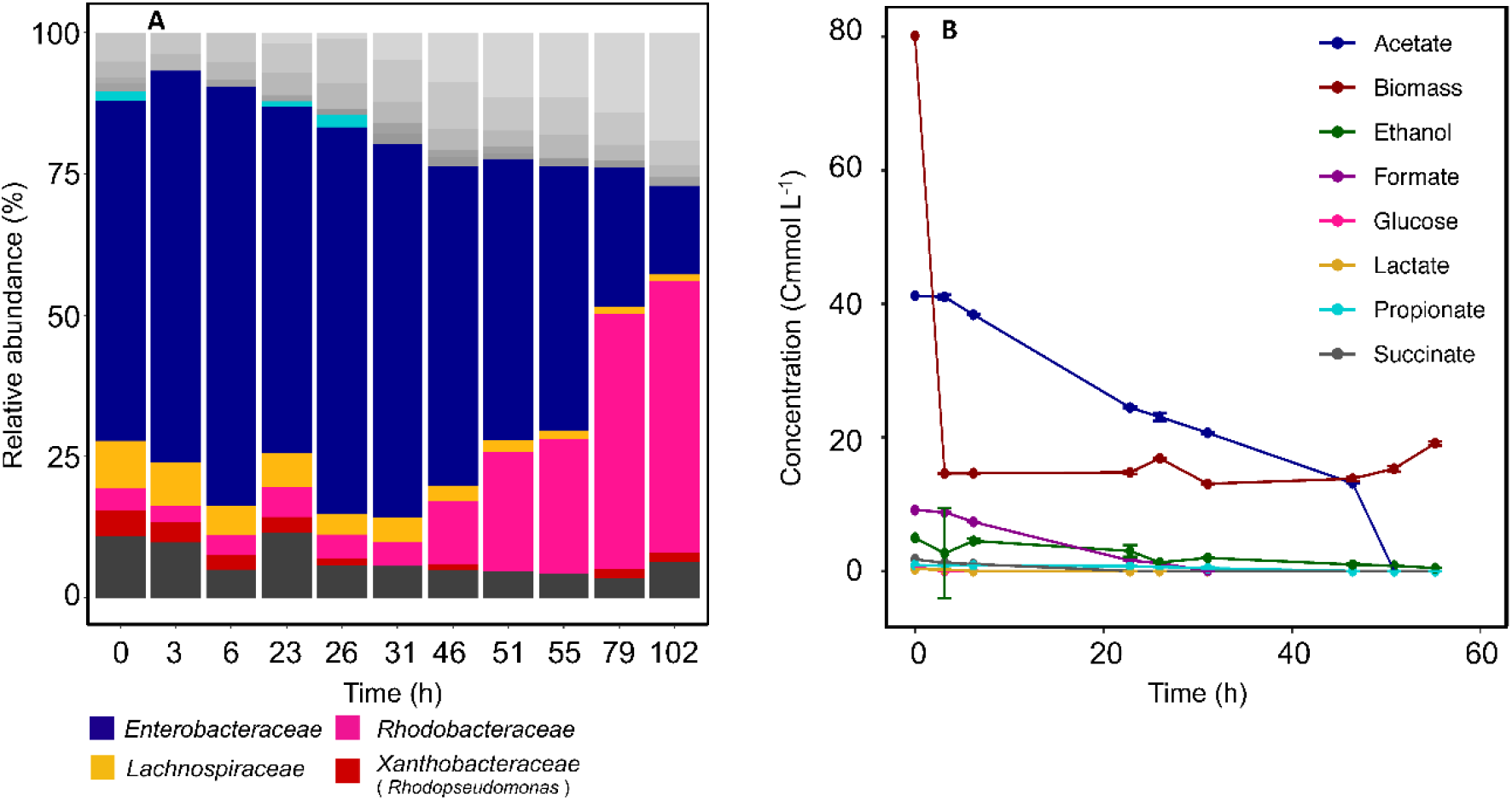
**A:** Microbial composition after the stop to the glucose feed. Over time, the FCB relative abundance decreased over time, while the *Rhodobacteraceae* increased. **B:** Fermentation products concentration after the glucose feed-stop. After 55 h all the fermentation products were depleted.

### Outlook

#### Metabolic strategies for microbial enrichment

Microorganisms use a combination of adaptive reactions to coexist with other organisms (Panikov, 2010). Based on the adaptation techniques, two major types of organisms can be identified (Andrews & Harris, 1986; Cowan et al., 2000): r- and K-strategists. The r-strategists show high growth and conversion rates, in low-populated and resources-rich environments. K-strategists thrive in highly populated environments, where the resources are limited. They exhibit lower growth rates but high substrate conversion yields.

Under the different reactor regimes, the microbial communities followed this postulate. *Enterobacteriaceae* were predominant under non-limiting conditions can be classified as r-organisms. *Clostridiaceae* and *Lachnospiraceae* showed high growth rates in nutrient-limiting environments and can be classified as K-strategists. Among the PPB guild, *Rhodopseuodmonas* showed a high-rate behaviour, dominating in substrate-rich environments. *Rhodobacter* showed high efficiency typical of the K-organisms, being able to survive also to nutrient-limited environments, as the CSTRs (**Table 4**).

**Table 4.** Reported maximum growth rates of the main FCB and PPB in the systems under glucose and acetate feed and adaptation strategies.

| Substrates | Maximum growth rate $\mu_{\max}$ (h<sup>-1</sup>) | | | | |
| --- | --- | --- | --- | --- | --- |
|  | Fermentative chemoorganoheterotrophic bacteria |  |  | Purple photoorganoheterotrophic bacteria |  |
|  | (FCB) |  |  | (PPB) |  |
|  | *Enterobacteriaceae* | *Lachnospiraceae* | *Clostridiaceae* | *Rhodopseudomonas* | *Rhodobacter* |
| Glucose | 0.3 – 1<br>(Hasona et al., 2004;<br>Khanna et al., 2012) | 0.3<br>(Soto-Martin et<br>al., 2020) | 0.25<br>(Gomez-Flores<br>et al., 2015) | 0.014<br>(Conrad & Schlegel,<br>1978) | 0.1<br>(Imam et al., 2011) |
| Acetate | - | - | - | 0.15<br>(Cerruti et al., 2020) | 0.05<br>(Kopf & Newman,<br>2012) |
| Selection<br>strategy | r-strategist<br>(on glucose) | K-strategist<br>(on glucose) | K-strategist<br>(on glucose) | r-strategist<br>(on acetate) | K-strategist<br>(on acetate) |

Kinetic parameters define the microbial selection mechanisms in CSTRs (Kuenen, 2019). In the CSTRs, the dilution rate was relatively low (0.04 h^-1^) compared to the growth rates of the fermentative organisms (0.5-1 h^-1^), as it was set to retain PPB in the system. In a continuous culture, the substrate (in this case glucose) is the limiting compound, and the organisms with higher affinity for it will be predominant (Andrews & Harris, 1986). Within the guild of fermentative organisms, *Clostridiaceae* and *Lachnospiaraceae* are expected to have a lower affinity constant (K_s_) and therefore have established over *Enterobacteriaceae*. The controlled pH and an appropriate dilution factor (D = 0.04 h^-1^) allowed the persistence of PPB. In particular, *Rhodobacter* was enriched in the glucose-fed CSTRs regardless the illumination conditions. We propose here a syntrophic association of FCB and PPB for glucose conversion toward CO_2_ and biomass (**Figure**). Glucose was first converted into acetate, ethanol, formate and lactate, and lactate was further transformed into acetate and propionate. All the fermentation products can be potentially used by PPB for growth.

#### Syntrophic associations between FCB and PPB

The interaction between fermentative organisms and purple bacteria is an example of syntrophic interaction. Syntrophy has been shown in biofilms and sludges, where a mass transport of compounds occurs between organisms (Rodríguez et al., 2006). Here, we showed the establishment of a metabolic cooperativity that lead to a multispecies consortium, combining sugar degradation with VFAs assimilation. In nature, PPB are present in many aquatic environments where IR light is present, and they have been found also in anthropogenic environments such as WWTPs. They harness their ability to grow on a diverse range of compounds, creating a wide metabolic net of interaction with other organisms.

The syntrophy between FO and PPB is not only important at an ecological level but can also be exploited for wastewater treatment. The FCB are used to treat food and agricultural wastes, to degrade sugars and produce biofuels as bioethanol and hydrogen gas of VFAs (Li et al., 2015; Thapa et al., 2015, 2019). Further efforts have been put to selectively direct PPB bioproduct formation toward one or another metabolic route (Puyol et al., 2017), with success. PPB, thanks to high biomass yields (Alloul et al., 2018), harbor a major industrial potential for the production of antioxidants and single-cell proteins. It has been proven that their biomass can be used as feed for shrimps without protein extraction (Qin et al., 2018).

#### Advantages of single-sludge vs. two-sludge processes for conversions of glucose by FCB and PPB

The single-step bioprocess implemented here has the advantage to simultaneously treat carbohydrate-rich wastewaters, factually removing the organic pollutants, and enriching organisms with high industrial potential, by combining the metabolic properties of FCB and PPB in one single sludge. An integrated single-stage process combining fermentation and purple photoorganoheterotrophy would allow to treat carbohydrate-rich wastewater in only one tank. The illumination conditions did not affect the syntrophy between FCB and PPB. A discontinuous illumination would reduce the operational costs compared to a continuously irradiated system (Qin et al., 2018). Understanding interactions between the FCB and PPB microbial guilds will help design efficient processes for carbohydrates-based wastewater treatment and valorization. If the aim is then to maximize and valorize the PPB biomass, a two-sludge process can be efficient to first ferment carbohydrates in the first tank selecting for FCB and then supplying the fermentation products like VFAs (whose spectrum can be engineered by mastering selection conditions) to the PPB tank.

## Conclusions

FCB and PPB interact as syntrophic organisms. FCB degrade glucose into VFAs, lactate and ethanol. Acetate was assimilated into biomass by PPB. The operational conditions are crucial to establish an interaction between the guilds. FCB are more efficient fermenters than PPB (although their metabolic versatility can allow for fermentation). Under glucose-fed batch regime FCB were enriched over PPB. Under glucose-fed CSTR conditions PPB population got enriched up to 30% in syntrophic association with FCB. An appropriate dilution rate (0.04 h^-1^) and pH regulation to 7 enabled to enrich PPB in glucose-rich environments by enabling their metabolic coupling with FCB.

## ACKNOWLEDGEMENTS

This study was funded by the start-up grant of the TU Delft Department of Biotechnology (Prof. David Weissbrodt). The authors warmly acknowledge Dirk Geerts, Zita van der Krogt and Ben Abbas for their technical assistance in fermentation and molecular biology labs at TU Delft.

## CONFLICT OF INTEREST STATEMENT

The authors share no conflict of interest.

## Notes

### Competing Interest Statement

The authors have declared no competing interest.

